# Diverse Genotype-by-Weather Interactions in Switchgrass

**DOI:** 10.1101/2021.08.19.456975

**Authors:** Alice H. MacQueen, Li Zhang, Samuel Pattillo Smith, Jason E. Bonnette, Arvid R. Boe, Philip A. Fay, Felix B. Fritschi, David B. Lowry, Robert B. Mitchell, Francis M. Rouquette, Yanqi Wu, Arbel Harpak, Thomas E. Juenger

## Abstract

The timing of vegetative and reproductive growth in plants (“phenological timings”) depend on genetic effects (G), environmental (e.g., weather) cues, and their interaction. Here, we measure phenological timings in two highly divergent switchgrass (*Panicum virgatum*) subpopulations using repeated plantings of cloned individuals at eight sites across the central United States. The timing of vegetative growth for the two subpopulations reversed between their two natural ranges and had strong negative correlations between these regions; in contrast, the timing of flowering was positively correlated between gardens. We expect that these phenotypic correlations consist of polygenic effects on phenology which have distinct patterns of GxE segregating at different mapped loci. Thus, we infer the mixture of ways genetic effects impact phenological timings, such as across common gardens (GxE) or with weather cues (GxWeather). We demonstrate that we can identify genetic variation with GxWeather and assign genetic loci to specific weather-based cues or other patterns. For example, in the Gulf subpopulation, 65% of genetic effects on the timing of vegetative growth covary with daylength 14 days prior to green-up date, and 33% of genetic effects on the timing of flowering covary with cumulative rainfall in the week prior to flowering. However, most variation in genetic effects cannot be attributed to variation in weather variables. Selective breeding for particular alleles at GxWeather loci could alter flowering responsiveness in a photoperiod or rainfall-specific way. More broadly, our approach refines the characterization of genotype-by-environment interactions and can be implemented in any species phenotyped in multiple environments.

## Introduction

Plant phenological traits are important components of plant fitness that are affected by multiple external environmental cues (e.g. degree of winter chilling, day length, temperature, soil fertility, and water availability), which signal existing or upcoming growing conditions (1–4). Genetic responses to environmental cues determine the speed, timing, and energy apportioned to vegetative and reproductive growth and shape plant physiological responses, lifespan, and lifetime production of viable seed. Day length (or photoperiod) is one of the most predictable environmental cues, and genetic sensitivity to photoperiod protects plants from potentially fatal consequences of phenological responses to temperature cues at the “wrong” time of year. However, the utility of specific environmental cues depends on both features of the environment, such as cue predictability and relevance, and the species’ adaptive strategies (5). Species with wide natural distributions can have multiple distinct environmentally cued phenological responses. For example, populations of sunflower (*Helianthus annuus*) exhibit day-neutral, facultative short day, and facultative long-day flowering responses, which vary with their environments (6, 7). These distinct genetic responses in different environments are known as genotype-by-environment interactions, or GxE.

Flowering time, or the transition from vegetative to reproductive growth, is a common subject of GxE research (6–13), a key output of selection driving adaptation to local environments (2, 14–17), and a selection target for crop improvement to adapt crops to local or future environments (18). Changing flowering responsiveness to photoperiod cues has allowed geographic range expansion and increased yields in several cereal species (14, 19–23) and other crops (24, 25). Recent statistical advances in studying phenological GxE have involved determining critical environmental indices before the phenological event occurs, such as photothermal time within a critical growth window (11). However, most studies of flowering GxE focus on finding a single, best fitting form of genotype-environment covariance, despite the key expectation that different genetic subpopulations, and even different genomic regions, have likely evolved distinct patterns of GxE (26).

Additionally, despite theoretical predictions that local adaptation should involve trade-offs caused by antagonistic pleiotropy, or alleles with effects with opposing fitness outcomes (27–30), previous empirical work has found limited evidence of trade-offs caused by this form of GxE (15, 31, 32). However, this work has been limited by a known statistical bias that reduced detection of genetic effects that differ in sign (31, 33, 34). Thus, despite substantial interest in the frequencies of various forms of GxE, the frequency of sign-changing GxE relative to other forms of GxE remains unknown.

Previous research suggests that switchgrass phenological timings should have GxWeather and that these timings could differ by genetic subpopulation. Switchgrass is considered a short-day plant with reproductive development strongly linked to day of the year (35). However, as part of its wide environmental adaptation across the eastern half of North America, its photoperiodicity has been predicted to differ by plant latitude of origin (36, 37). We previously found that divergent Midwest and Gulf genetic subpopulations of switchgrass have distinct sets of environmental adaptations associated with each of two fitness proxies, biomass and overwinter survival (38). The Midwest genetic subpopulation is primarily composed of individuals from the well-studied upland switchgrass ecotype (39, 40), while the Gulf subpopulation has individuals from the well-studied lowland ecotype and the phenotypically intermediate coastal ecotype (38).

Here, we test for GxWeather for the timing of two phenological traits by loading patterns of genetic effects on phenology at eight common gardens onto many patterns of weather covariance at these gardens. To do this, we phenotyped a diversity panel of hundreds of switchgrass genotypes from the Midwest and Gulf subpopulations for the timing of vegetative and reproductive development. We did this at eight common garden locations spanning 17 degrees of latitude: these gardens covered the majority of the latitudinal and climatic range of switchgrass and captured the most comprehensive picture to date of the environmental variation this species encounters. We defined multiple ways phenological traits might covary with weather (Table 1) and additional ways phenological traits might vary by garden (SI Appendix, Section S1), then jointly re-estimated genetic effects on these timings at all eight common gardens using the set of these covariance matrices that significantly improved the modeled log-likelihood when included (SI Appendix, Section S2-S5) (41). We used the Bayesian framework *mash* (multivariate adaptive shrinkage) developed by (41), to refine effect size estimates from genome-wide association (GWAS). *Mash* allowed us to identify and specify multiple covariance structures among genetic effect estimates across sites, including structures that represent covariance in weather variables of interest. Importantly, this method circumvented statistical biases in detecting genetic effects with the same or opposite signs (42). To confirm our genetic mapping of GxWeather, we compared genomic locations of the significant posterior effect estimates from *mash* to mapping results from an outbred mapping population grown at the same sites. Our analyses allowed us to describe the weather cues and types of GxE affecting phenology in two divergent natural populations of switchgrass across the species’ latitudinal range.

**Table 1.** Weather variables and time frames prior to the start of vegetative and reproductive growth used to construct the hypothesis-based covariance matrices. Covariance matrices were constructed for, and tested on, three subpopulations and two phenological traits. The correlations between values of these weather variables for genetically identical plants grown in different gardens were used to fill off-diagonal cells of the covariance matrices. Narrow-sense heritabilities for these values at each garden were used for the diagonal cells.

| Weather variable | Time Frame | Covariance Matrices |
| --- | --- | --- |
| cumulative growing degree days at 12C in the time frame | (1-7), 14, 21, 28 days prior to the phenological date | 60 |
| cumulative rainfall in the time frame | (1-7), 14, 21, 28 days prior to the phenological date | 60 |
| day length (hours) on a specific day indicated by the time frame | (1-7) and 14 days prior to the phenological date | 48 |
| day length change (seconds) on a specific day indicated by the time frame | (1-7) and 14 days prior to the phenological date | 48 |
| average temperature in the time frame | (1-7), 14, 21, 28 days prior to the phenological date | 60 |

## Results

Genotypes from the Gulf and Midwest subpopulations had distinct phenological trait timings and distinct patterns of phenological trait correlations across our eight common garden sites (Figure 1). At the three Texas common gardens (hereafter ‘Texas’ gardens), located within the natural range of the Gulf subpopulation, the onset of vegetative growth, or ‘green-up date’, for Gulf genotypes occurred before Midwestern green-up, and the onset of reproductive growth, or ‘flowering date’, for Gulf genotypes occurred after Midwestern flowering (Figure 1 A). At the four northernmost common gardens (hereafter ‘North’ gardens), located within the natural range of the Midwest subpopulation, both green-up and flowering of Gulf genotypes occurred after that of Midwest genotypes. At the Oklahoma common garden, located near the natural range limits of both the Gulf and the Midwest subpopulations, Gulf and Midwest green-up occurred over the same period, and Gulf genotype flowering occurred after Midwestern flowering (Figure 1 A). These patterns led to strong negative phenotypic correlations for the onset of vegetative growth between the North and Texas gardens, particularly in the Gulf and in the population containing all individuals from the Gulf and Midwest subpopulations (hereafter, ‘Both’ subpopulations) and contributed to positive phenotypic correlations for the onset of flowering that had larger magnitudes at more northern gardens (Figure 1 B).

**Figure 1.**
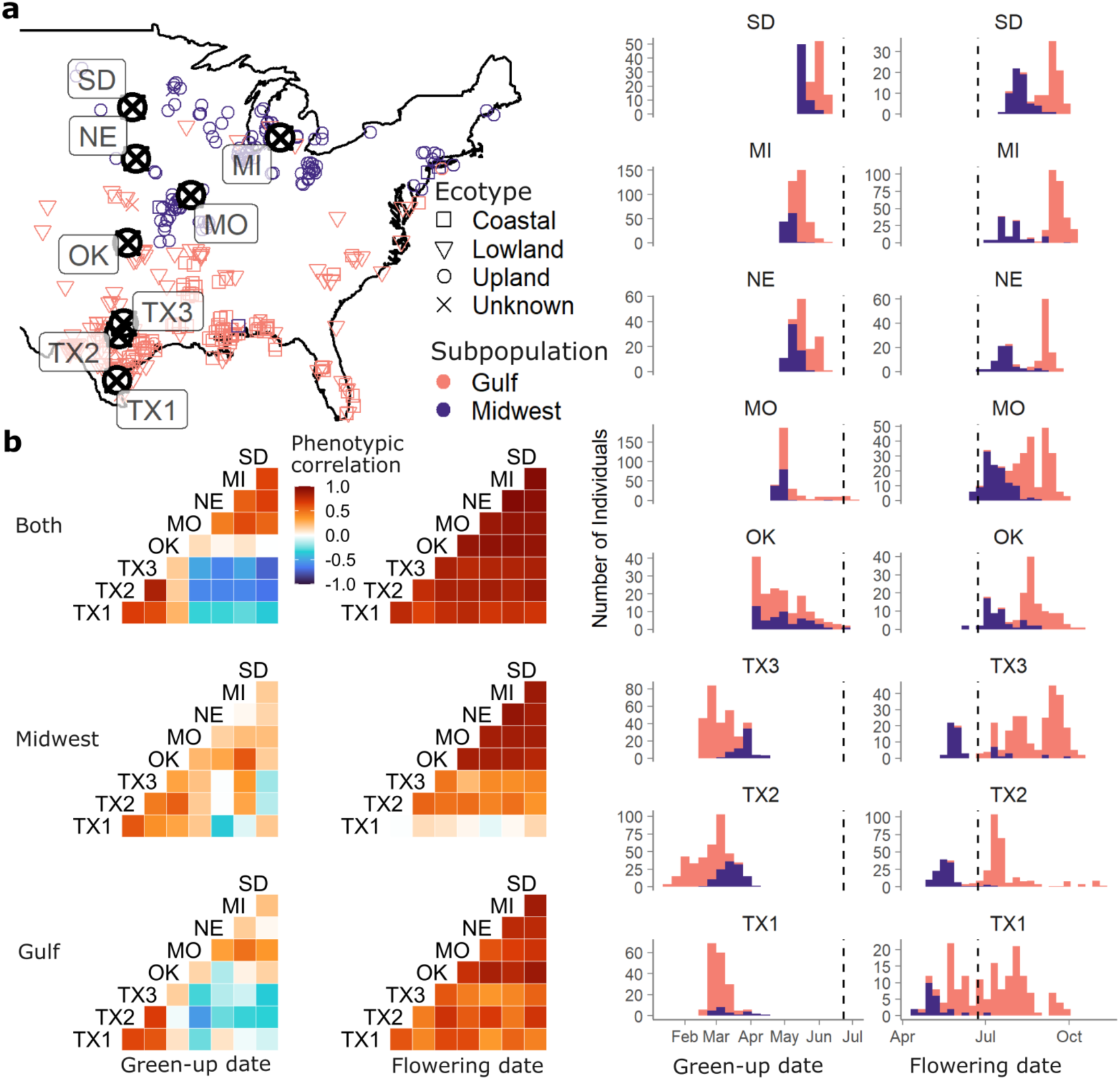
Characterization of the timing of the onset of vegetative (green-up) and reproductive (flowering) growth in the switchgrass diversity panel. (a) Map and trait histograms of green-up and flowering dates across two genetically distinct switchgrass subpopulations and eight common gardens. Purple represents individuals from the Midwest genetic subpopulation, and pink individuals from the Gulf subpopulation; map positions represent the original collection locations for the genotypes, and shapes represent the ecotype of the genotype. Histogram vertical dashed lines indicate the summer solstice. Common gardens are arranged in latitudinal order. (b) Phenotypic correlations between clonal replicates planted at eight common gardens, within and between two genetic subpopulations.

Narrow-sense heritabilities (h^2^) suggested that rank-changing GxE for these phenotypes was present across the common gardens (SI Appendix, Figure S1). h^2^ were typically high at individual gardens: 59% on average for green-up date, and 87% for flowering date. However, h^2^ was variable across gardens, and green-up dates were uncorrelated (r^2^ < 0.2) or negatively correlated between pairs of gardens (Figure 1 B). These negative and small correlations undoubtedly contributed to the low h^2^ values for green-up and flowering date when estimated jointly at all eight gardens: h^2^ was 0.8% for green-up and 23.2% for flowering date (SI Appendix, Figure S1).

### Inference of genome-wide patterns of GxE and GxWeather for vegetative and reproductive timing

We expected patterns of GxE and GxWeather to vary at the locus, subpopulation, and trait level. Thus, we were interested in modeling the different ways polygenic effects might covary across our eight common gardens(43). We jointly re-estimated the genetic effects of a subset of SNPs across all eight common gardens using *mash* models, which can flexibly capture many modes of covariance across gardens (SI Appendix, Sections S1-S5, Datasets 1-6). To obtain genetic effects, we used subsets of effects from garden- and subpopulation GWAS (SI Appendix, Section S2-S3; Table S1). To obtain covariance structures, we specified both covariance structures introduced in the initial *mash* manuscript (e.g., garden-specific effects; equal effects at all gardens, Figure 2 A), and GxWeather covariance matrices estimated from the covariance of empirical weather patterns at each garden at specific times before the phenological event (Table 1; SI Appendix, Section S1). Then, we used a greedy algorithm to select covariance matrices from each model’s set that significantly improved the model likelihood (SI Appendix, Section S4; Table S2). Finally, we fit *mash* models using the selected set of covariance matrices on a subset of relatively unlinked (r^2^ < 0.2), ‘strong’ genetic effects with low *p*-values (SI Appendix, Section S5).

**Figure 2.**
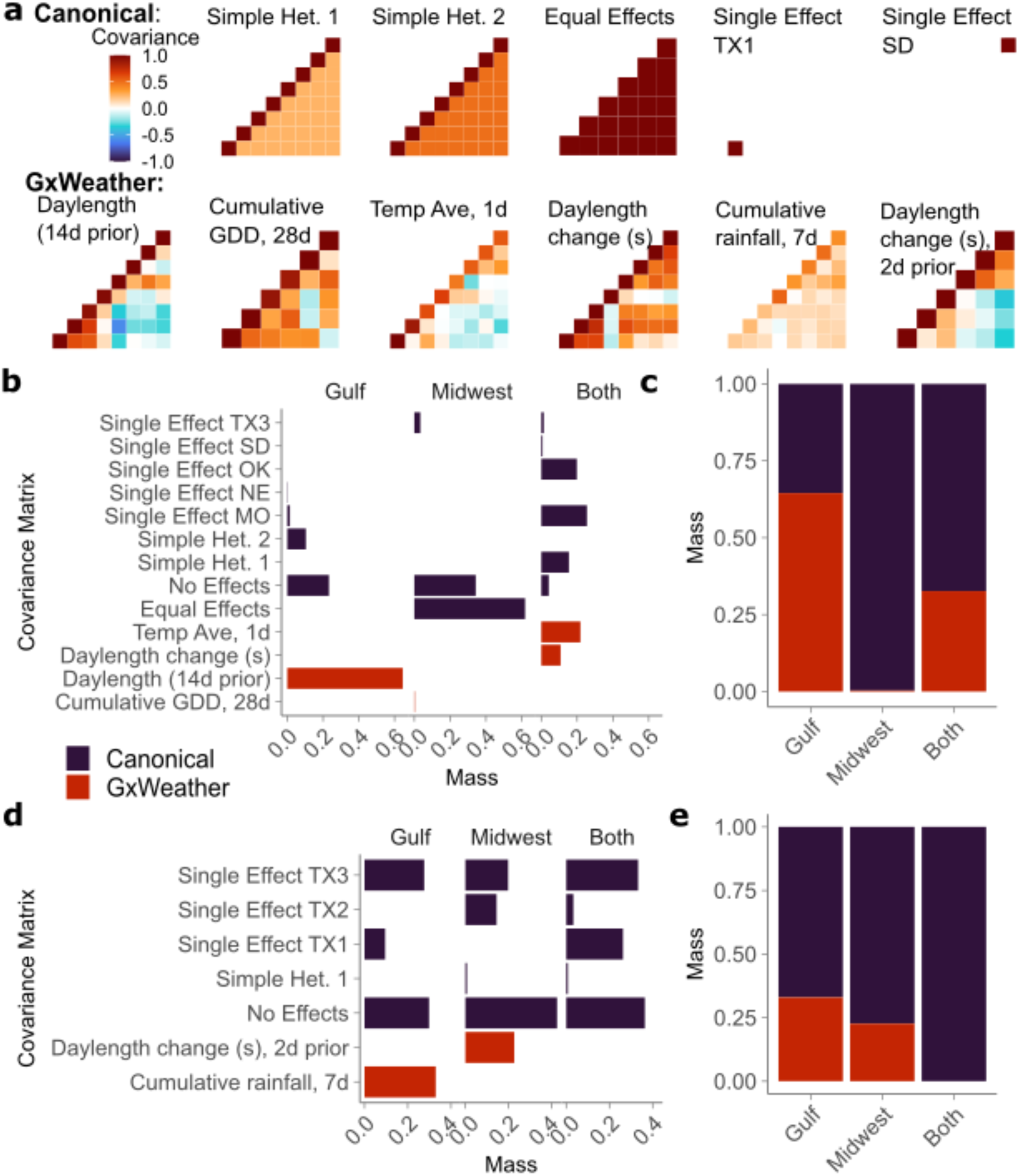
Example Canonical and GxWeather covariance matrices specified in *mash* and the posterior weights placed on all covariance matrices. (a) Common gardens are arranged in latitudinal order within the matrices. Top row: Four example canonical covariance matrices. Canonical matrices (purple) have simple interpretations, such as equal effects across all common gardens, or effects specific to a single common garden. Bottom row: Five example GxWeather covariance matrices specified for the green-up date or flowering date phenotype; these matrices were created from environment-specific correlations across eight common gardens, and are described in Table 1. (b,d) Total posterior weight placed on each covariance matrix type specified for (b) green-up date and (d) flowering date *mash* models, within and between two genetic subpopulations. Covariance matrices included in *mash* that had zero posterior weight in all three *mash* runs on the genetic subpopulations, such as the identity matrix, are not shown. (c,e) Total posterior weight placed on covariance matrices that were GxWeather or Canonical, for the (c) green-up date phenotype and (e) flowering date phenotype.

The loadings of genetic effects onto the GxE and GxWeather covariance matrices included in each model provide information on genome-wide patterns of SNP-environment interaction. For example, the phenotypic correlations for the onset of vegetative growth had moderate negative correlations between the Texas and North gardens, particularly in the Gulf subpopulation and Both subpopulations (Figure 1 B). If these phenotypic correlations have a genetic component, they could be partially or completely captured by the covariance structures specified in the *mash* model. In this case, the joint estimate of effects for many SNPs could have high mixture proportions, or mass, on a covariance matrix that captures the negative correlation between Texas (TX1, TX2, and TX3) and North (MO, NE, MI, and SD) gardens. In our data, daylength 14 days prior to green-up date was negatively correlated between Texas and North gardens (Figure 2 A); if many SNP effects have mass on this GxWeather matrix, we would infer that the effects of those SNPs have that form of GxWeather covariation.

Nine GxWeather covariance structures were selected by the greedy algorithm, one to two per subpopulation: six for green-up date, and three for flowering date (SI Appendix, Table S2). Of these nine, six had mass (>0.1%) on them in *mash* models of the strong effects (Figure 2 B,D). Five of these six matrices had negative covariances between gardens in the Texas and North regions (Figure 2 A), while one had all positive or near-zero covariances.

For green-up date, SNP-associated phenotypic effects covaried with different weather-based cues in the Gulf & in Both subpopulations (Figure 2 B). In total, 65% of the posterior weight of strong SNP effects in the *mash* model of Gulf green-up fell on a covariance matrix constructed using the covariance of day length 14 days prior to the date of green-up. The covariance matrix for this weather cue was visually similar to the observed pattern of phenotypic correlation for the timing of vegetative growth in the Gulf subpopulation (Figure 1 B; Figure 2 A). *Mash* models of the timing of vegetative growth in the Midwest subpopulation and Both subpopulations did not include this GxWeather covariance. The Midwest had 0.15% weight on a GxWeather covariance matrix of cumulative GDD for the 28d prior to green-up, a GxWeather matrix that was visually similar to the Midwest’s observed pattern of phenotypic correlation (Figure 1 B). Both subpopulations had non-zero weights on two additional GxWeather covariance types, average temperature one day prior to green-up, and the day length change in the day prior to green-up (Figure 2 B). The average temperature covariance matrix had negative covariances between Texas and North gardens, though not as strong as the negative phenotypic correlations observed in Both subpopulations (Figure 1 B; Figure 2 A). The Gulf subpopulation and Both subpopulations had substantially more mass on GxWeather covariance matrices than the Midwest population (Figure 2 C) for the onset of vegetative growth.

For flowering date, distinct GxWeather covariance structures captured covariance in effect sizes for SNPs in the Gulf and Midwest subpopulations. 33% of SNP effects on flowering in the Gulf subpopulation covaried with cumulative rainfall in the seven days prior to flowering (Figure 2 D). 22.6% of SNP effects on flowering in the Midwest subpopulation covaried with day length change in the two days prior to flowering (Figure 2 D), which had negative covariances between Texas and North gardens. Neither of these covariance matrices significantly improved the log-likelihood of the *mash* model of Both subpopulations, and no GxWeather covariance matrices had non-zero mass in models of Both subpopulations (Figure 2 E). In five of the six *mash* models of strong effects, the GxWeather covariance matrices captured a minority of the posterior weights of the strong effects (Figure 2 C,E); in these five models, the majority of this mass was on various canonical covariance matrices. These matrices included simple heterozygosity, with intermediate, positive covariances between all gardens, and single effect matrices with garden-specific effects. Re-estimation of genetic effects with *mash* allowed us to identify and quantify the fraction of loci exhibiting GxWeather patterns; a minority of GxE was GxWeather covariance, and the majority of GxE covariance in these models is driven by other, unknown drivers of GxE.

### Frequency of rank-changing GxE in significant SNP effects

We expected to observe common rank-changing GxE at the level of individual loci as this is a key theoretical prediction of local adaptation (27–30). Previous empirical work has found limited evidence of trade-offs caused by antagonistic pleiotropy; however, this work had a known statistical bias reducing the detection of effects that differed in sign (15, 31, 32). To determine the frequency of rank-changing GxE, we used the local false sign rate (lfsr), an analogue of the local false discovery rate that establishes confidence in the effect sign, not the effect’s difference from zero, to determine significance. We required lfsr significance (p < 0.05) in both gardens to include effects. This means that our tests for a sign change between gardens carry an equal statistical burden to those for effects with the same sign. We separated kinds of effects at the level of individual loci into SNP effects that differ in sign between gardens (effects with rank-changing GxE between gardens), SNP effects that differ in magnitude (effects that are large in one garden or region and smaller in others), and SNP effects that are indistinguishable in two gardens (similarly large or small in comparisons between gardens) (Figure 3 A).

**Figure 3.**
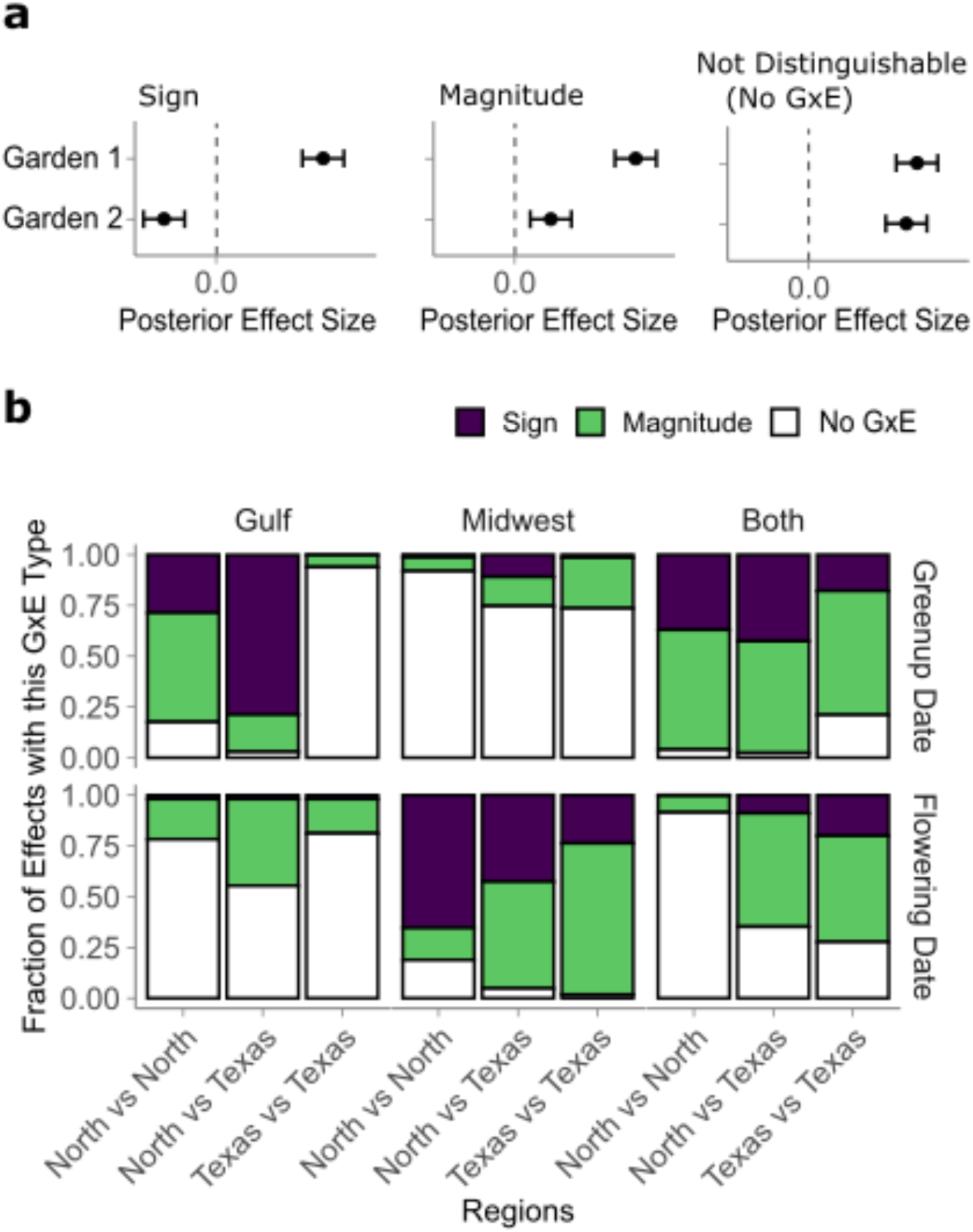
Types of GxE present between of pairs of jointly re-estimated SNP effects in eight common gardens, for effects with lfsr < 0.05 at both gardens for each pair of gardens contrasted. a) Examples of effect patterns at three pairs of sites with three types of GxE. All effects are for an alternate allele, with the reference allele effect defined to be zero at both gardens and represented by the dashed vertical line. Sign: Effects that differ in sign at these pairs of gardens (p < 0.05, lfsr). Magnitude: Effects identical in sign (p < 0.05, lfsr) that differ in magnitude by a factor of >0.4. Not Distinguishable: Effects not distinguishable by magnitude nor sign of the effect, with no measurable GxE. Confidence intervals are illustrative that the effect estimate does not overlap zero. b) The fraction of effects with each GxE type for the onset of vegetative growth (green-up date) and reproductive growth (flowering date), within and between two genetic subpopulations. Common gardens are grouped by the larger region they came from: North gardens are within the natural range of the Midwest subpopulation, and include MO, NE, MI, and SD, while Texas gardens are within the natural range of the Gulf subpopulation, and include TX1, TX2, and TX3.

For green-up date for the Gulf subpopulation, hundreds to thousands of pairwise effects exhibited rank-changing GxE, or a difference in sign between pairs of Texas and North gardens (Figure 3 B; SI Appendix, Fig S2). 78.7% of pairwise comparisons between North and Texas gardens had a difference in sign, while only 28.6% and 0.2% of North-North or Texas-Texas comparisons had a difference in sign, respectively (Figure 3 B). The majority of pairwise effects for greenup for the Midwest (>73%; 68-342 effects) were indistinguishable, or had no GxE. The majority of pairwise effects for Both subpopulations (>85%) were the same sign, and effects most often differed in magnitude between gardens within and between the regions; however, these differences in effect sign and magnitude were mostly between the MO site and other sites (Figure 3 B; SI Appendix, Fig S2).

For flowering date for the Gulf subpopulation, less than 2% of pairwise effects exhibited rank-changing GxE, or a difference in effect sign within or between regions (Figure 3 B). More effects differed in magnitude between the Texas and North regions than within these regions (42.7% vs <20%; Figure 3 B), while the majority of effects had no GxE. The Midwest population had relatively few significant effects for flowering (0-174 per pairwise comparison), but a large proportion of these differed in sign between Texas and North regions (42.7%) or within the North region (65.4%). Finally, in Both subpopulations, less than 20% of pairwise effects differed in sign (Figure 3 B). Most differences in sign were between TX1, the southernmost garden, and all other gardens (SI Appendix, Figure S3). Similarly, more effects that differed in magnitude included gardens in the Texas region (52.3-55.9%), and most effect pairs in the North region were not distinguishable (91.5%).

### Confirmation of effects on phenology using an independent mapping population

We sought additional experimental support the occurrence and location of our significant SNP effect re-estimates using an independent pseudo-F2 mapping population created from Gulf & Midwest individuals and grown at the same sites (Figure 4 A,B). We conducted quantitative trait loci (QTL) mapping of flowering as functions of five environmental cues that we also used as covariance matrices in *mash*, and identified eight QTL for flowering date, eight QTL for flowering as a function of day length change two days prior, one QTL for the start of vegetative growth, and two QTL for vegetative growth as a function of daylength change one day prior, all of which showed QTL by environment interactions (SI Appendix, Figure S4). All QTL for flowering overlapped with one or more homologs from rice or *A. thaliana* with functionally validated roles in flowering (SI Appendix, Dataset 7); the QTL on Chr04K overlapped a gene recently functionally validated for flowering in switchgrass (44). All flowering and green-up QTL intervals contained at least one SNP significant in at least one *mash* run at a log10-transformed Bayes Factor > 2, or in the 1% tail of significance, whichever was stricter (SI Appendix, Dataset 8). We also looked for enrichments of *mash* SNPs in the 1% tail of significance (the ‘*mash* 1% tail’) within each QTL interval. At the 5% level, three QTL had enrichments of SNPs in the *mash* 1% tail. Overall, there were five significant enrichments (p < 0.05, hypergeometric test) of SNPs in the *mash* 1% tail in the QTL intervals. Thus, we were able to experimentally support the genomic windows of some re-estimates of significant SNP effects with a QTL mapping experiment using a separate mapping population.

**Figure 4.**
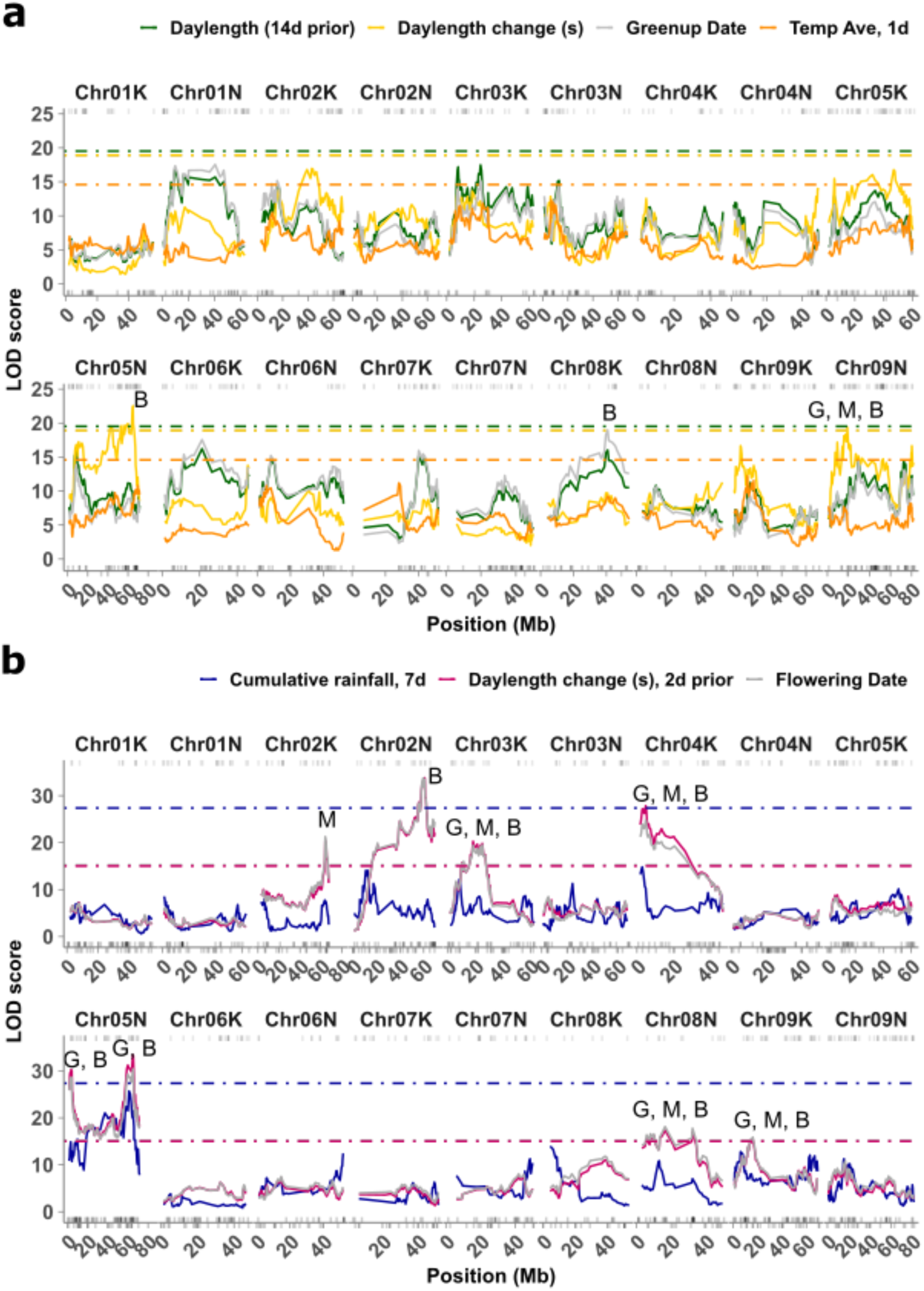
Overlaps of QTL from an outbred pseudo-F2 cross and with jointly re-estimated SNP effects in the 1% tail of significance from a diversity panel. Dotted lines indicate permutation-based significance thresholds for each weather-related function. Stars indicate QTL with significant enrichment for SNPs in the 1% *mash* tail; G, M, and B indicate which subpopulation had enrichment: G - Gulf subpopulation, M-Midwest subpopulation, B - both subpopulations. Rug plots show genomic locations of SNPs in the 1% *mash* tail for flowering date for each subpopulation: Both subpopulations are above the plot panel, the Gulf subpopulation is above the x-axis, and the Midwest subpopulation is below the x-axis. a) QTL mapping for the onset of vegetative growth (green-up date), and three weather-related functions of green-up date. b) QTL mapping for the onset of reproductive growth (flowering date), and two weather-related functions of flowering date.

## Discussion

As the climate and the natural environment change, it is increasingly critical to understand how patterns of plant-environment interactions will change in response. To do this, we must understand the current patterns of trait covariation across environments, the genetic underpinnings of these patterns, and the cases where this covariation can be altered through selection at individual loci. Here, we demonstrate that we can associate multiple patterns of GxWeather with specific genomic regions using a switchgrass diversity panel grown at eight common gardens. We assigned genetic effects to both GxWeather patterns with interpretable weather-based cues, and to unmeasured, site-based patterns. We used this approach to study GxWeather for the timings of vegetative and reproductive development in the deeply genetically diverged Gulf and Midwest subpopulations of switchgrass.

Our analysis of the timing of vegetative and reproductive growth revealed many alleles with rank-changing GxE, or sign changes in their effects between the Texas and North gardens (Figure 3 B). The Gulf & Both subpopulations had rank-changing GxE for green-up date, while the Midwest subpopulation had rank-changing GxE for flowering date. As phenological timings are major components of plant fitness, this result supports theoretical models that local adaptation should involve trade-offs due to antagonistic pleiotropy at the level of individual loci (27–30). Experimental designs in local adaptation research now often use more than two field sites and a wide range of genetic variation; *mash* and the local false sign rate facilitate the analysis of these experimental designs. We use *mash* to determine that rank-changing GxE for phenological traits is common in small genomic regions (15, 45, 46).

Our analysis of the timing of flowering showed that the Gulf and Midwest subpopulations have distinct GxWeather: flowering timing in the Midwest subpopulation has photoperiod-related genetic variation, in that flowering timing covaries with a day length change signal two days before flowering occurs. In contrast, the Gulf subpopulation does not have genetic variation in flowering that covaries with a photoperiod cue. Instead, the Gulf subpopulation has genetic variation in flowering that covaries with the rainfall that occurs in the week prior to flowering. Three genomic regions affecting flowering that we re-estimated across all eight sites were also supported by QTL from an independent mapping population at these sites (Figure 4 B). Models combining both subpopulations showed less signal, perhaps due to the distinctness of cues in both subpopulations or to confounding caused by population structure.

Identifying the environmental cues that are predictive of, or even correlated with, plant phenotypic responses remains a major challenge to studies interrogating gene action across many natural environments. The GxWeather photoperiod and cumulative rainfall cues we identify here are functions of the genotypes measured and capture only a minority of SNP effects on flowering. We could only assign SNP effects to a GxWeather covariance structures in four of the six phenotype & genetic subpopulations we modeled. It’s likely we did not include some GxWeather important to these phenological cues - for example, overwintering parameters that might cause variation in the start of vegetative growth. More generally, it is difficult to predict the time scales over which individuals may integrate environmental cues, particularly in perennial species which may integrate these cues over longer time scales. If this integration time itself varies between individuals, the covariance structures we modeled cannot reelect this, though these structures would likely be highly correlated with GxWeather structures we did include. Our approach offers an opportunity to specify multiple environmental cues and compete them to explain patterns of genetic effects, allowing us to detect how important these cues are genome-wide, and how strongly each cue influences each SNP. This is a key development to further improve our understanding of genetic variation in GxE.

## Materials and Methods

Whenever possible, plant material will be shared upon request. Source data and code to replicate these analyses are available at: https://github.com/Alice-MacQueen/pvdiv-phenology-gxe.git. SNP data to replicate these analyses are available from the UT dataverse at https://doi.org/10.18738/T8/A604BU.

### Scoring onset of vegetative and reproductive development in two mapping panels

In 2019, we scored two phenological events every two days in two mapping populations of switchgrass, a diversity panel and a pseudo-F2 cross, planted at eight common garden locations (38, 40, 46). We scored the onset of vegetative growth, or green-up date, as the day of the year when 50% of the tiller area of the crown of the plant cut the previous year had green growth. The onset of reproductive growth, or flowering date, was the day of the year when 50% of the plant tillers had panicles undergoing anthesis.

The formation and resequencing of the diversity panel has been described previously (38). The diversity panel contained 134 sequenced, clonally propagated individuals from the Midwest genetic subpopulation, and 229 from the Gulf genetic subpopulation. To allow for the possibility that different subpopulations had different strengths of connection between our phenotypes and genotypes (47), we conducted three sets of genetic analyses: on Gulf and Midwest genotypes separately, and on both subpopulations together (‘Both’ subpopulations). Analyses to determine narrow-sense heritability (h^2^) for green-up and flowering were done using linear mixed models and followed (38) (SI Appendix, Section S6).

To confirm candidate genomic regions found using *mash* on the diversity panel, we analyzed flowering in an outbred pseudo-F2 cross between four individuals, two Midwest and two Gulf individuals. The formation of this mapping population has been described previously (40); additional details on QTL mapping can be found in SI Appendix, Section S7. To be directly comparable to the diversity panel data, only 2019 phenology data from the pseudo-F2 cross from the same eight common garden sites were used.

### Joint re-estimation of SNP effects to assess the frequency of rank-changing GxE and assignment of genome-wide patterns of GxE and GxWeather

We were interested in specifying genetic models for trait variation that allowed more than one form of GxE, as we reasoned that different loci should display different forms of GxE (e.g., changes in effect magnitude vs sign). In addition, we were interested in an unbiased estimation of the frequency of rank-changing GxE relative to other forms of GxE, as the presence of rank-changing GxE at the level of individual loci is a key theoretical prediction of local adaptation. *Mash* allowed us to both specify multiple forms of GxE and GxWeather and conduct unbiased statistical tests for when SNP effects changed sign between common gardens.

To use *mash* on our diversity panel, we had to specify both a relatively uncorrelated set of covariance matrices, which in our case defined types of GxE and GxWeather between gardens, and we had to specify subsets of SNP effect estimates and standard errors for our traits at each common garden. To specify a set of covariance matrices, we first defined many covariance matrices, including GxWeather matrices that represented the correlation in weather cues between gardens before the phenological event (SI Appendix, Section S1), then implemented a model selection approach that used a greedy algorithm to evaluate if the log likelihood of the *mash* model was significantly improved as additional covariance matrices were included (SI Appendix, Section S4). To specify subsets of SNP effect estimates and standard errors to use for both the greedy algorithm and with the optimal set of covariance matrices, we first calculated best linear unbiased predictors (BLUPs) for each phenological trait in each genetic subpopulation and each common garden (SI Appendix, Section S2). Next, we determined effect estimates for 8.8 to 12.3 million SNPs per subpopulation by conducting garden-specific GWAS on these BLUPs using *k* vectors of singular values to correct for population structure, where *k* was the smallest integer value that made the genomic control coefficient closest to 1 (SI Appendix, Section S2). Singular values were computed using singular value decomposition of the matrix of all SNPs, with iterative SNP pruning and removal of regions in long-range linkage disequilibrium (48). Third, to make the *mash* models computationally feasible, we extracted two subsets of SNP effect estimates and standard errors from our GWAS effect estimates: (i) effects from a subset of “strong” tests corresponding to stronger effects on our traits; (ii) results from a random subset of all tests to correspond to an unbiased representation of all effects (SI Appendix, Section S3). We used the subset of random effects in our greedy *mash* algorithm (SI Appendix, Section S4) and used the subset of strong effects in our *mash* models with the optimal set of covariance matrices (SI Appendix, Section S5).

The loadings of genetic effects onto the multiple covariance structures specified in our *mash* models provided information on genome-wide patterns of SNP-environment interaction. In addition, the the GxWeather covariance structures allowed hypothesis testing of specific weather variables as cues for the start of vegetative and reproductive growth. Say that our diversity panel contains a SNP in the gene CONSTANS (CO), a well-known flowering time regulator, and that only one of the alleles affects the promotion of flowering in a photoperiod-dependent manner. In that case, the joint estimate of effects for that SNP could have a high mixture proportion, or mass, on a covariance matrix created using a photoperiod-based environmental cue, such as day length at some interval prior to flowering. In our data, we would infer that the effect of that SNP on flowering was cued by the weather variable used to create the GxWeather covariance structure.

Our joint re-estimation of SNP effects also allowed us to characterize the overall patterns of GxE in the set of SNPs where there was pairwise significance of effects at pairs of gardens. To do this, we used the ‘get_GxE’ function of the switchgrassGWAS R package. First, this function determines the set of SNPs with evidence of significant effects in both conditions for all pairs of conditions using local false sign rates (lfsr) as the significance criteria. Then, this function determines if effects significant in both conditions are of opposite sign.

Using the lfsr rather than the local false discovery rate (local FDR) is a critical change in our ability to detect alleles directly contributing to rank-changing GxE between environments. The local FDR, like other measures of FDR, focuses on if we have enough evidence to reject the null hypothesis that an effect j is 0, or that there is a significant effect. Previous studies of antagonistic pleiotropy (e.g. (46)) have used the local FDR or equivalent statistical tests to detect antagonistic pleiotropy. These tests were conservative, in that they required two non-zero effects of different signs, while tests for differential sensitivity required only one non-zero effect. This previous work recognized that this testing bias could lead to undercounting occurrences of antagonistic pleiotropy (31, 34), and sought to reduce it by permutation (33). However, using the lfsr to test for allelic effects that differ in sign does not undercount these occurrences, as this statistic answers a fundamentally different question. For each effect *j*, the *l fsr_j_* is defined as the probability that we make an error in the sign of effect *j* if we were forced to declare the effect positive or negative (42). Thus, rather than asking “Are these two effects different?” - as we reasonably expect two effects to be, even if this difference cannot be measured - the local false sign rate answers a more meaningful question: Can we be confident in the sign of this effect?

In addition, the get_GxE function also sets an arbitrary threshold to count an effect as changing in magnitude between environments, commonly known as differential sensitivity or a change in amplitude of the effect. For differential sensitivity, this function determines if effects significant in both conditions are of the same sign and of a magnitude (not tested for significance) that differs by a factor of 0.4 or more. The remaining effects that are significant in both conditions have the same effect sign and similar effect magnitudes and we denote these effects as having no GxE. The distinction between effects with different magnitudes is arbitrary but useful to fully characterize how effects vary across environments to ultimately influence phenotypes. Our use of the lfsr to determine significance and our specification that SNP effects must be significant in both conditions to be included means that our tests for alleles with rank-changing GxE carry an equal statistical burden to those measuring differential sensitivity and effects without GxE.

## Acknowledgements

We thank the Brackenridge Field laboratory, the Ladybird Johnson Wildflower Center, and the Juenger laboratory for support with plant care and propagation. The authors acknowledge the Texas Advanced Computing Center (TACC) at The University of Texas at Austin for providing HPC storage resources that have contributed to the research results reported within this paper. This material is based upon work supported in part by the Great Lakes Bioenergy Research Center, U.S. Department of Energy, Office of Science, Office of Biological and Environmental Research under Award Numbers DE-SC0018409 and DE-FC02-07ER64494, the US Department of Energy Awards DE-SC0021126 and DESC0014156 to T.E.J., DE-SC0017883 to D.B.L, NIH grant R35GM151108 and a Pew Scholarship to A.H., National Science Foundation PGRP Awards IOS0922457 and IOS1444533 to T.E.J, and the Long-term Ecological Research Program (DEB 1832042) at the Kellogg Biological Station.

