## Supplementary material for "Diverse Genotype-by-Weather Interactions in Switchgrass": SI Appendix

PNAS

#### This PDF file includes:

Figs. S1 to S4
Tables S1 to S2
Legends for Dataset S1 to S8
SI References

#### Other supporting materials for this manuscript include the following:

Datasets S1 to S8

### Supplementary Methods

**Section S1. Generation of covariance matrices to provide multiple ways SNP effects could covary across common gardens.** A trait measured in two environments can be considered two traits that are genetically correlated (Falconer 1952). By extension, a phenological trait measured at eight common gardens can be modeled as eight genetically correlated traits represented by a covariance matrix, where diagonal entries represent the additive genetic variance present in the population at that garden, and non-diagonal entries represent the genetic correlation between the trait in a pair of environments.

We were interested in the types of genetic correlation that could be expressed by different alleles across the genome and reasoned that different loci could have different patterns of genetic correlation. For example, the effects of an allele of locus $A$ could covary with a weather-based cue if $A$ had a common allele, $a$, that was responsive to this cue. In contrast, the effects of a different locus $B$ might have an effect at only one garden, for example if a pathogen was only present at one garden, and $B$ has a common allele $b$ that is resistant to that pathogen.

We specified three qualitative categories of covariance structures across gardens: "canonical" covariance, with simple patterns of effect size covariance introduced in the initial *mash* manuscript; "data-driven" covariances derived from common patterns of SNP effects observed in the data and introduced in the initial *mash* manuscript, and "GxWeather" covariance, estimated from the covariance of empirical weather patterns at each garden over specific time frames before each genotype underwent the phenological event.

Canonical covariance matrices were created by *mash* and fell into four groups: an identity matrix, an equal effects matrix, singleton matrices, and simple heterozygousity matrices. The identity matrix has values of zero for covariance between different gardens and correlations of one within gardens, and represents the situation in which the effects in different conditions are independent. The equal effects matrix has values of one for all matrix elements, and represents identical effects across all conditions. Each singleton matrix has a value of one for a single garden and zeros for all other elements, representing a garden-specific effect. Three simple heterozygousity matrices are generated, which have covariances of 0.25, 0.5, or 0.75 for all off-diagonal matrix elements, and correlations of one for diagonal entries, representing moderately positively correlated effects across all sites.

Data-driven covariance matrices can also be created by *mash* and fall into two groups: an overall pattern of covariance between all conditions, denoted as $tPCA$, and the first five principle components of a singular value decomposition of the $tPCA$, denoted as $PCA_1$ through $PCA_5$ (Urbut et al. 2018).

To create the GxWeather covariance matrices we transformed each genotype's green-up and flowering date at each garden to create date-related values that were summary functions for a weather variable across a specified date range. For example, the cumulative sum of the rainfall that occurred seven days prior to the flowering date was used to create one possible GxWeather cue variable for flowering date. The code for these calculations can be found at git@github.com:Alice-MacQueen/pvdiv-phenology-gxe/R/Analysis_v1.2_mash_using_greedy_covar.qmd, starting at line 387.

We defined the two phenological dates as functions of five weather variables and eight to ten time frames (Table 1). First, cumulative growing degree days (GDD) variables for each time frame were calculated as $GDD = \sum_{DATE-i}^{DATE} max(T_{mean} - T_{base}, 0)$, where $T_{mean}$ is the daily average temperature, defined as $(T_{max} + T_{min})/2$, $T_{base}$ is the base temperature of 12°C for switchgrass, $T_{max}$ is the maximum daily temperature, $T_{min}$ is the minimum daily temperature, $DATE$ is the phenological date, and $i$ is the time frame, or the number of days prior to the phenological date that GDD is summed across (Kiniry et al. 2005; Behrman et al. 2013). Second, cumulative rainfall values in each time frame were calculated as $C_{rain} = \sum_{DATE-i}^{DATE} PRCP$, where PRCP is the daily rainfall measured in millimeters, $DATE$ is the phenological date, and $i$ is the time frame, or the number of days prior to the phenological date that rainfall is summed across. Third, day length in hours was determined for each time frame as a specific single day prior to the phenological date, as indicated by the time frame $i$, e.g. $DATE - i$. Day length was calculated as a function of latitude and day of the year as in (Forsythe et al. 1995). Fourth, day length change in seconds was determined for a specific single day prior to the phenological date as indicated by the time frame, as $DYLN_{change(s)} = (DYLN_{DATE-i} - DYLN_{DATE-i-1}) * 60 * 60$, where $DYLN$ in the day length in hours, $DATE$ is the phenological date, and $i$ is the number of days prior to the phenological date. Fifth, average temperature in each time frame was defined as $T_{ave} = \sum_{DATE-i}^{DATE}(T_{max} + T_{min})/2$, where $T_{max}$ is the maximum daily temperature, $T_{min}$ is the minimum daily temperature, $DATE$ is the phenological date, and $i$ is the time frame, or the number of days prior to the phenological date that average temperature is summed across.

We then generated GxWeather covariance matrices derived from correlations in these GxWeather cue variables for the three subpopulations. These covariance matrices represent the correlations between genotypes for these weather-related variables across our eight common gardens. To create these matrices, we determined the correlations between these weather-related cue variables for identical genotypes grown at different common gardens and used these correlations to fill the off-diagonals of a eight-by-eight matrix. To fill the diagonal entries of these matrices, we used the narrow-sense heritability for each weather-related cue variable at each garden, calculated using the same methods as Section S3. Matrices need to be positive semi-definite to be used in *mash*, so when the resultant matrices were not positive semi-definite, the `Matrix::nearPD` function was used to compute the nearest positive definite matrix to the correlation matrix (Bates, Maechler, and Jagan 2023). Finally, all covariance matrices were rescaled so that the maximum diagonal element of the rescaled matrix is 1, as in (Urbut et al. 2018). Code used to generate these matrices can be found at git@github.com:Alice-MacQueen/pvdiv-phenology-gxe.git/R/Analysis_v1.2_mash_using_greedy_covar.qmd, starting at line 715.

We defined distinct sets of GxWeather covariance matrices for the two phenological dates and for the three genetic subgroups. We used the same five weather variables to transform both green-up and flowering date, but defined the weather-based covariance matrices relative to each of the two phenological dates. GxWeather covariance matrices were also defined separately for individual genotypes from the Gulf, Midwest, and Both subpopulations.

**Section S2. Univariate genome-wide association to obtain initial effect estimates.** To obtain initial effect estimates for each single nucleotide polymorphism (SNP) for each combination of phenological trait, genetic subpopulation, and common garden, we conducted univarate genome-wide association (GWAS) on site-specific best linear unbiased predictors (BLUPs) using the switchgrassGWAS package and the methods in (Lovell et al. 2021). The full SNP diversity panel generated in (Lovell et al. 2021) was subset within each subpopulation for all analyses to retain only SNPs with $\leq$ 20% missing data and minor allele frequencies $>$ 0.05. This subsetting resulted in 8.8 million SNPs retained for the Midwest subpopulation, 10.3 million SNPs retained for the Gulf subpopulation, and 12.3 million SNPs retained for Both subpopulations.

BLUPs were calculated independently for each combination of trait, genetic subpopulation, and common garden. We used ASReml to estimate BLUPs using mixed models of the form:

$$\mathbf{y} = 1 + Zu + e$$

$$Var(u) = G\sigma_u^2$$

$$Var(e) = I\sigma_e^2$$

where the vector $\mathbf{y}$ represents the flowering date or greenup date values for that garden, $Z$ the design matrix for random effects, $u$ the whole genome additive genetic effect, $e$ the residual, $G$ the genomic relationship matrix based on all SNPs retained for subpopulation-specific analyses, and $I$ the rank-$\mathbf{y}$ identity matrix (Clark and Werf 2013). ASReml can specify considerably more complex variance structures; however, we were interested in specifying covariance structures using *mash*, which allows the specification of multiple covariance structures; hence, we used a relatively simple model to calculate BLUPs for use in univariate GWAS. The code for these calculations can be found at git@github.com:Alice-MacQueen/pvdiv-phenology-gxe.git/R/Analysis_v1.2_mash_using_greedy_covar.qmd, starting at line 69.

Univariate GWAS were run independently for each combination of trait, genetic subpopulation, and common garden. The code used to run GWAS can be found at git@github.com:Alice-MacQueen/pvdiv-phenology-gxe.git/R/Analysis_v1.2_mash_using_greedy_covar.qmd, starting at line 240. We used `pvdiv_standard_run()`, an opinionated function of the switchgrassGWAS R package, to conduct GWAS. To control for population structure and run GWAS, this function uses a three-part process. First, a fast truncated singular value decomposition (SVD) is generated from the SNP data, which has initial pruning of regions in long-range linkage disequilibrium (LD) and iterative pruning and removal of regions in long-range LD (Privé et al. 2018). By default, 15 vectors of singular values, $k$, are computed. Pruning and removal of long-range LD regions are recommended in research in human genetics to best control for population structure (Abdellaoui et al. 2013; Privé et al. 2018; Price et al. 2008). Second, linear regression models using 0-15 of these vectors of singular values were used to run GWAS and to calculate the genomic control coefficient, $\lambda_{GC}$, for each $k$ (Devlin and Roeder 1999). Third, we chose the lowest value of $k$ that made $\lambda_{GC}$ closest to 1, and ran GWAS using a linear regression model with this $k$ for population structure correction.

At this stage, we reviewed the univariate GWAS results, both Manhattan and QQ-plots, for population structure issues that were not resolved using the $k$ that minimized $\lambda_{GC}$. These issues were apparent in the data in two ways: (i) univariate GWAS with many significant SNPs with extremely similar $p$-values, usually indicative of genetic effects that, by chance or, more likely, because of confounding effects of population structure, were identical across many SNPs and thus not resolvable to a single genomic location; (ii) univariate GWAS with many effect estimates and standard errors of zero, indicating an inability to estimate effects given the constraints of small population size & population structure. We only noticed these issues in our smallest subpopulation, the Midwest subpopulation, in the green-up date trait at the MO and SD garden, and in the flowering date trait at the OK and NE gardens. Due to these issues, we dropped the MO and SD garden from subsequent green-up analyses for the Midwest subpopulation, and we dropped the OK and NE gardens from subsequent Midwest flowering date analyses.

**Section S3. Selection of effect estimate subsets to model using *mash*.** To make the subsequent analyses using *mash* computationally feasible, we followed the author's recommended analysis outline for large datasets using *mash* and considered subsets of effects at different stages of the mash analysis. We extracted two subsets of tests from our complete GWAS effect estimate datasets: 1) effects from a subset of "strong" tests corresponding to stronger effects on our traits; 2) results from a random subset of all tests to correspond to an unbiased representation of all effects. We also clumped effects before selection of both subsets so that signal strength was standardized across different LD blocks in the genome, and not limited to closely linked associations.

The intuition behind taking a subset of relatively unlinked, randomly selected effects is that this will capture a realistic fraction of the null and non-null effects in the full dataset, and thus be an unbiased representation of the amount and types of signal in the full dataset. Mash then uses this subset of random effects to estimate the number of null effects in the dataset and shrink some estimates towards null effects, effectively, towards a covariance matrix with no effects. The intuition behind using a subset of relatively unlinked effects with strong effects, or low $p$-values, is that these effects are the most likely to end up with significant jointly re-estimated SNP effects, and every SNP in a linkage block is likely to have effects re-estimated similarly.

To prepare subsets of effect estimates and their standard errors from these GWAS for use in mash, we used `pvdiv_bigsnp2mashr()` from the switchgrassGWAS R package. For each phenological trait and each genetic subpopulation, we created two subsets of effects and standard errors in paired $m \times n$ matrices, where $m$ were marker effects and $n$ were common gardens. The first subset of effects and standard errors, the strong effects, was created by selecting relatively unlinked ($r^2 < 0.2$) SNPs from each univariate GWAS that had the smallest $p$-values in those GWAS. Notably, the majority of these "strong" effects were not significant at a 5% level in univariate GWAS. The number of SNPs from each univariate GWAS was chosen to result in approximately million entry matrices for each trait & subpopulation. We selected 19K - 33K effects and standard errors for the strong effect subsets (Table S1). In many cases, there was overlap in the markers with the largest effect estimates at different locations; thus, 19 - 33K SNPs selected for each of eight univariate GWAS ultimately gave sets of 35K - 63K SNPs (Table S1). The second subset of effects and standard errors, the random subset, was created by selecting relatively unlinked ($r^2 < 0.2$) SNPs from each univariate GWAS after clumping using minor allele frequency. To account for the chance that true effects could be sparse, we doubled the size of the random subset relative to the strong subset. We selected 38 - 66K SNP effects and formed subsets of 71 - 127K SNP markers for the random subset.

**Section S4. Greedy *mash* algorithm to select covariance matrices that significantly improved the model log-likelihood.** Because we were particularly interested in determining whether genomic loci varied in their patterns of genetic covariance across gardens, and thus loaded on to different covariance matrices in *mash*, specifying the optimal set of covariance matrices in *mash* was an important model-building problem. If very similar covariance matrices are included when fitting a model in *mash*, *mash* will arbitrarily assign effect loadings across these matrices, as all assignments will lead to the same fit to the effect-size data. Our GxWeather covariance matrices were in many cases highly correlated, as they represented covariation in the same weather variables in overlapping time frames. We thus evaluated the performance of *mash* models when different covariance matrices were included by implementing a "greedy" mash algorithm.

Specifically, given $k$ covariance matrices for each phenotype, and $m$ corresponding effect sizes and standard errors across $n$ contexts of interest, we first fit a set of mash models with each of the $k$ matrices separately. We then identified the model with the maximum log likelihood estimates and selected the corresponding matrix, leaving $k-1$ covariance matrices. We then fit a new set of mash models where the selected covariance matrix is paired with each of the remaining $k-1$ covariance matrices. The most likely pair of covariance matrices was then selected and the process was repeated for all possible $k-2$ matrix triplets. This process was repeated until one of two stop conditions was met: (i) all covariance matrices have been added to the mash model, resulting in a final model with all of the original $k$ covariance matrices are included or (ii) a likelihood ratio test with one degree of freedom between the likelihood of the current model and the model from the previous iteration has a $p$-value $>$ 0.05. The result is a set of log likelihood estimates for each model and the stepwise 'path' of covariance matrices that best explain the observed effect sizes and standard errors observed for the phenotype of interest.

We applied the greedy *mash* algorithm to the entire set of canonical, data-driven, and GxWeather covariance matrices described in Section S1. We used the random subset of SNP effect estimates, described in Section S3, in the greedy *mash* algorithm in order to test covariance matrices on an unbiased representation of all effects across the genome. The code to replicate this analysis can be found at git@github.com:Alice-MacQueen/pvdiv-phenology-gxe.git/analysis/mash_greedy_algorithm/mash_h2_diag_covar/ in four directories: all_inputs/, gulf_inputs/, midwest_flowering/, and midwest_greenup/. For our six models of two phenological traits in three genetic subpopulations, between eight and twelve iterations of the greedy algorithm were necessary before a stop condition was met (Table S2). The relatively small number of necessary iterations, and correspondingly small number of covariance matrices selected, indicate that the hypothesis covariance matrices were highly correlated with one another and that inclusion of all the hypothesis covariance matrices would introduce redundancy into the model. Most notably, the canonical "Identity" covariance matrix, representing independent effects, was the first matrix selected in all six applications of the greedy algorithm (Table S2). This indicated that a large number of independent effects were present in our analyses of both phenological traits. In addition, most or all of the available canonical covariance matrices, which specifed garden-specific effects, were selected before any GxWeather covariance matrices or data-driven covariance matrices (Table S2). Finally, at least one GxWeather covariance matrix was selected in each greedy algorithm application; in contrast, one data-driven covariance matrix was selected in just one of the six models (Table S2).

**Section S5. *Mash* to jointly re-estimate SNP effects across eight common gardens.** To evaluate the prevalence and kinds of covariance patterns of SNP effects across our eight common gardens, we used multivariate adaptive shrinkage (*mash*) to jointly re-estimate SNP effect estimates for our strong effects subset using the set of covariance matrices selected by the greedy *mash* algorithm. *Mash* is a statistical method that allows estimation and comparison of many effects jointly across many different conditions; it improves on previous methods by allowing arbitrary correlations in effect sizes among conditions. *Mash* assigns mixture proportions for each SNP's effect onto each provided covariance matrix using maximum likelihood. Then, *mash* uses Bayes' theorem to shrink effects for each SNP towards the set of covariance matrices in accordance to their mixture proportions. The ability to specify multiple types of covariance matrix is an important advantage *mash* offers for studying patterns of GxE: user-specified covariance matrices allow hypothesis testing of specific drivers for each SNP's effect, such as weather-related cues, while the data-driven covariance matrices *mash* creates allow exploration of additional unexplained patterns of covariation.

The code used to run *mash* can be found at git@github.com:Alice-MacQueen/pvdiv-phenology-gxe.git/R/Analysis_v1.2_mash_using_greedy_covar.qmd, starting at line 1340.

**Section S6. Narrow-sense heritability estimation.** In the diversity panel, we determined narrow-sense heritabilities ($h^2$) for green-up and flowering dates at single gardens using genomic relationship matrices calculated using the van Raden method (VanRaden 2008). Genomic relationship matrices were calculated within each subpopulation (Midwest and Gulf) and for both genetic subpopulations (Both). We used rrBLUP (Endelman 2011) to specify mixed models of the form:

$$\mathbf{y} = 1 + Zu + e$$

$$Var(u) = G\sigma_u^2$$

$$Var(e) = I\sigma_e^2$$

in which the vector **y** represents the flowering date or green-up date values for that garden, Z the design matrix for random effects, u the whole genome additive genetic effect, and e the residual. Matrix G is the whole genomic relationship matrix based on all SNPs retained for subpopulation-specific analyses. I is the rank-y identity matrix. Phenotypic variance $\sigma_p^2$ is $\sigma_u^2 + \sigma_e^2$. Narrow-sense heritability is then $h^2 = (\sigma_u^2/\sigma_p^2)$.

These models were run for each of the eight gardens, and across all gardens by adding an additional environmental effect of site without an interaction term. This resulted in 54 models: 3 sets of populations (the Gulf, Midwest, and Both subpopulations) for 9 garden sets (all eight gardens separately, and all eight gardens together) and two phenotypes (green-up date and flowering date).

**Section S7. Outbred pseudo-F2 mapping population and Quantitative Trait Locus mapping.** To confirm candidate genomic regions and patterns of allelic effects found in the diversity panel, we analyzed flowering in an outbred pseudo-F2 cross between four grandparents, two Midwest and two Gulf individuals. The formation of this mapping population has been described previously (Milano, Lowry, and Juenger 2016). The parents of this cross were DAC, an early flowering Midwest individual, VS16, a late flowering Midwest individual, AP13, an early flowering Gulf individual, and WBC, a late flowering Gulf individual. We made F1 crosses of the two early flowering genotypes, AP13xDAC, and the two late flowering genotypes, WBCxVS16. We then clonally propagated and planted the four parents, the two F1 genotypes (AP13xDAC, and VS16xWBC), and 801 F2 genotypes at eight field sites in May-July of 2015. To be directly comparable to the diversity panel data, only 2019 phenology data from the pseudo-F2 cross from the same eight common garden sites were used here.

Details on the genetic map construction, map polishing and fine-scale reordering can be accessed on DataDryad. QTL mapping was conducted with R/qtl2 (Broman et al. 2019). We performed a genome scan with a linear mixed model that accounts for the relationships among individuals and for environmental covariates (i.e., field sites). The full model can be expressed as:

$$phenotype = \mu + QTL + E + QTL * E + kinship + e$$

where $\mu$ is the population mean, QTL is the marker genetic effect, E is the environmental effects (here, common garden), QTL*E is the interaction between marker genetic and environmental effects, kinship corresponds to the background polygenic variation, and e is the error term. The genome scan was accomplished with the 'scan1' function. The statistical significance of the genome scan was established by performing a stratified (i.e., stratifying on common garden) permutation test (n=1000) using 'scan1perm' function. The estimated QTL effects (Figure S4) were obtained using 'scan1coef' function in R/qtl2.

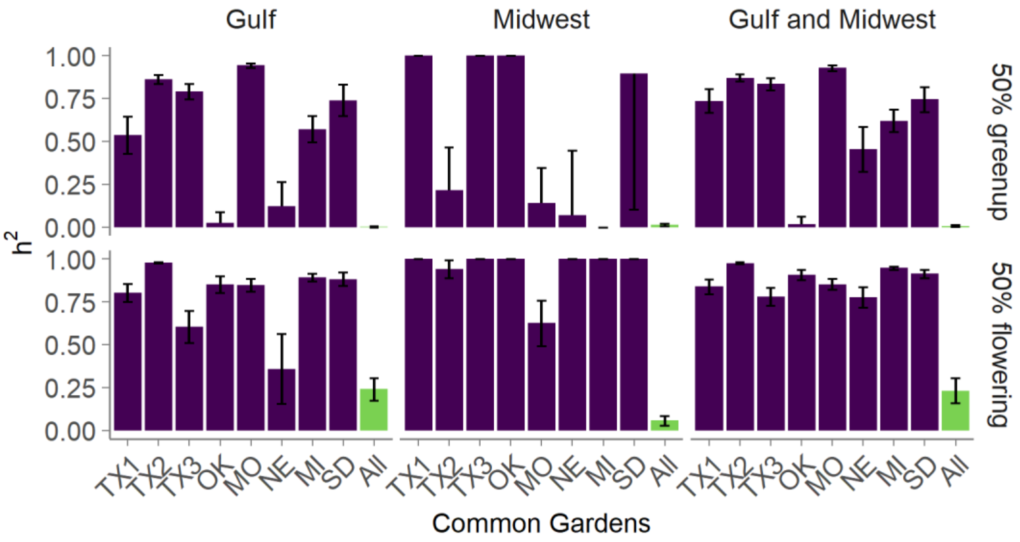


**Figure S1.** Narrow sense heritability of green-up date and flowering date within single common gardens (purple) and across all eight common gardens (green), within and between two genetic subpopulations.

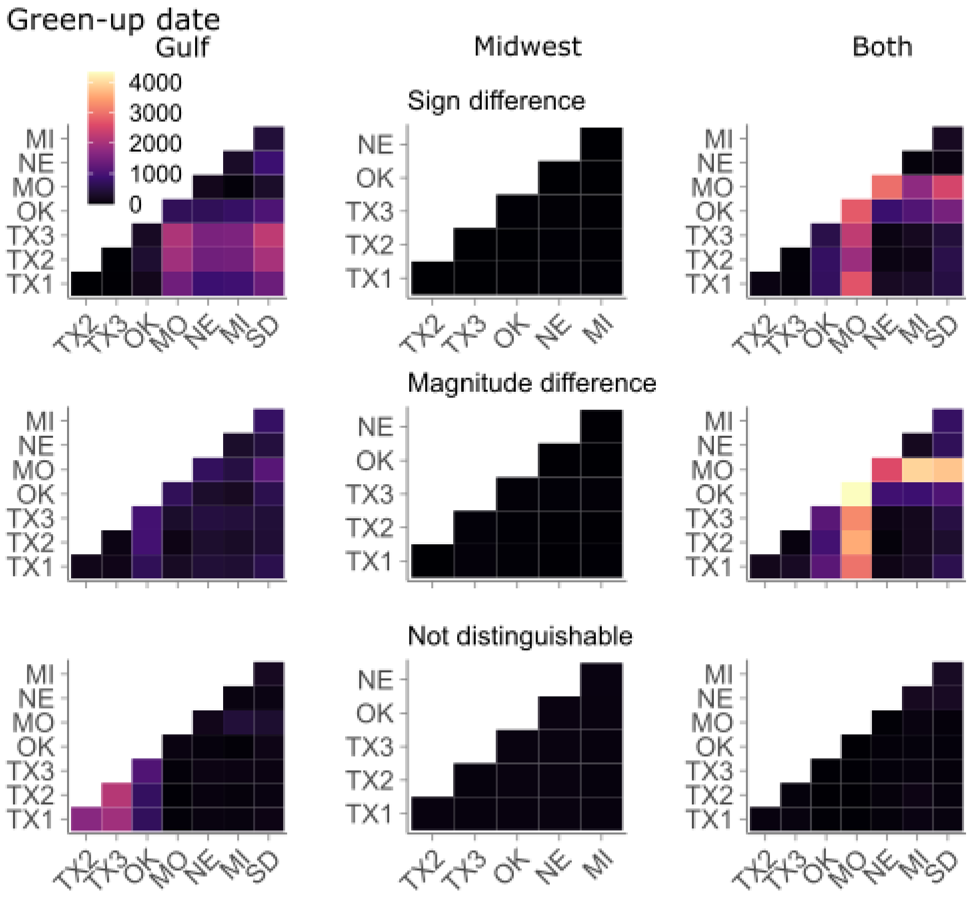


**Figure S2.** Types of GxE present between of pairs of jointly re-estimated effects in eight common gardens, for effects with lfsr < 0.05 at both gardens in each pair. Effect patterns: Sign: Effects that differ in sign at these pairs of gardens (p < 0.05, lfsr). Magnitude: Effects identical in sign (p < 0.05, lfsr) that differ in magnitude by a factor of >0.4. Not Distinguishable: Effects not distinguishable by magnitude nor sign of the effect, with no measurable GxE. b) The number of effects with each GxE type for the onset of vegetative growth (green-up date) within and between two genetic subpopulations.

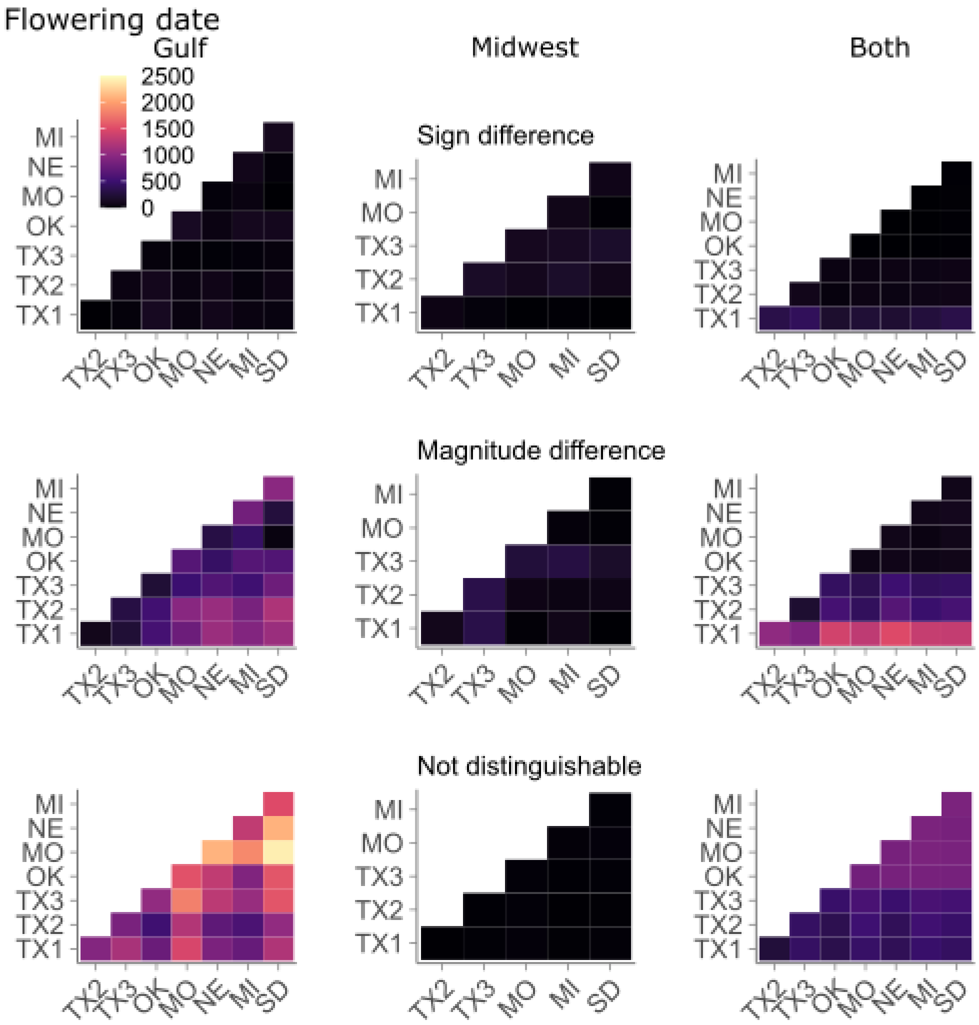


**Figure S3.** Types of GxE present between of pairs of jointly re-estimated effects in eight common gardens, for effects with lfsr < 0.05 at both gardens in each pair. Effect patterns: Sign: Effects that differ in sign at these pairs of gardens ($p < 0.05$, lfsr). Magnitude: Effects identical in sign ($p < 0.05$, lfsr) that differ in magnitude by a factor of >0.4. Not Distinguishable: Effects not distinguishable by magnitude nor sign of the effect, with no measurable GxE. b) The number of effects with each GxE type for the onset of reproductive growth (green-up date) within and between two genetic subpopulations.

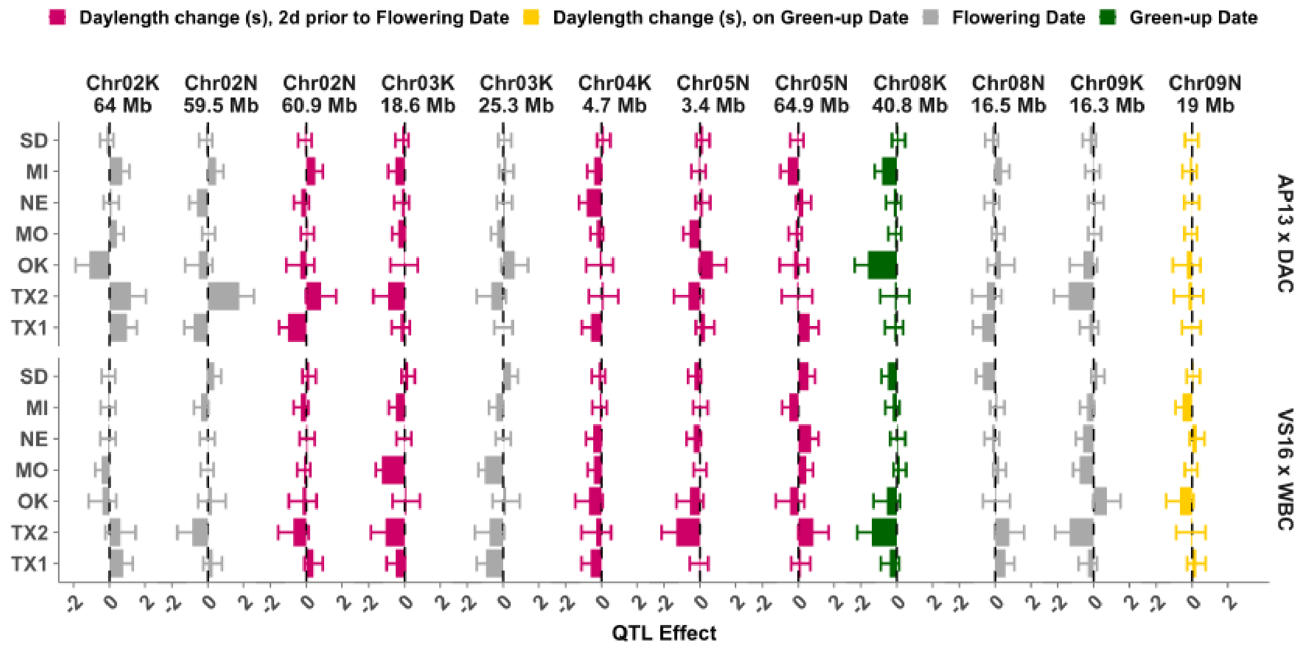


**Figure S4.** All QTL identified in the outbred mapping population displayed GxE, in that all QTL had effects that differed in at least two gardens. Gardens are on the y-axis in latitudinal order; the effects of the significant QTL in Figure 4 are displayed as differences between allelic effects for the two sets of grandparent lines in the outbred mapping panel.

**Table S1. Effect estimates selected for joint re-estimation across eight common gardens using *mash***

| Phenology type | Subpopulation | Effects Selected per GWAS, Strong Subset | Effects Selected per GWAS, Random Subset | Total Markers, Strong Subset | Total Markers, Random Subset |
|---|---|---|---|---|---|
| Greenup Date | Gulf | 19000 | 38000 | 45931 | 91862 |
| | Midwest | 33000 | 66000 | 57729 | 115458 |
| | Both | 19000 | 38000 | 44765 | 89530 |
| Flowering Date | Gulf | 19000 | 38000 | 50490 | 100980 |
| | Midwest | 33000 | 66000 | 63367 | 126734 |
| | Both | 19000 | 38000 | 35735 | 71470 |

**Table S2. Covariance matrices selected in each iteration of the greedy *mash* algorithm, for each phenological trait and genetic subpopulation. Naming matches the conventions in Figure 2 of the main text.**

| Greedy Algorithm Iteration # | Gulf Greenup Date | Midwest Greenup Date | Both Greenup Date | Gulf Flowering Date | Midwest Flowering Date | Both Flowering Date |
|---|---|---|---|---|---|---|
| 1 | Identity | Identity | Identity | Identity | Identity | Identity |
| 2 | Simple Het. 2 | Single Effect OK | Single Effect MI | Simple Het. 1 | Simple Het. 1 | Single Effect TX3 |
| 3 | Single Effect MO | Equal Effects | Single Effect SD | Single Effect MI | Single Effect MI | Single Effect TX1 |
| 4 | Single Effect NE | Single Effect TX3 | Single Effect NE | Single Effect TX3 | Single Effect TX2 | Single Effect OK |
| 5 | Daylength (14d prior) | Single Effect TX2 | Single Effect TX2 | Single Effect SD | Single Effect MO | Single Effect MI |
| 6 | Single Effect MI | Cumulative GDD, 12C, 1d | Single Effect OK | Single Effect OK | Single Effect TX3 | Single Effect NE |
| 7 | Single Effect SD | Cumulative GDD, 12C, 28d | Single Effect TX3 | Single Effect NE | Daylength change (s), 2d prior | Single Effect TX2 |
| 8 | Single Effect TX2 | - | Simple Het. 1 | Single Effect TX1 | Single Effect SD | Single Effect MO |
| 9 | Cumulative GDD, 6d | - | Single Effect TX1 | Single Effect TX2 | - | Single Effect SD |
| 10 | Single Effect TX3 | - | Single Effect MO | Single Effect MO | - | Simple Het. 1 |
| 11 | Data-driven $PCA_2$ | - | Temp Ave, 1d | Cumulative rainfall, 7d | - | Cumulative rainfall, 1d |
| 12 | - | - | Daylength change (s) | - | - | - |

**Datasets.** Datasets 1 through 6 are csv files where the first two columns have the following definitions:

- Marker: The SNP marker in the format Chromosome_Position
- log10BF: log10(Bayes Factor) of the significance of the marker effect in the mash model

The remaining column names follow the pattern Effect_[Mean/StandardError/lfsr]_[Subpopulation]_[Phenotype]_[Garden], where Mean and Standard Error are estimates of the effect mean and standard error, lfsr is the local false sign rate statistic for the effect, and [Subpopulation], [Phenotype], and [Garden] follow the conventions of Figure 1.

**SI Dataset S1 (Dataset_1_Gulf_vegetative_growth_date.csv)**

SNP-associated effects for the start of vegetative growth jointly re-estimated in the Gulf genetic subpopulation.

**SI Dataset S2 (Dataset_2_Midwest_vegetative_growth_date.csv)**

SNP-associated effects and standard errors for the start of vegetative growth jointly re-estimated in the Midwest genetic subpopulation.

**SI Dataset S3 (Dataset_3_Both_subpopulations_vegetative_growth_date.csv)**

SNP-associated effects and standard errors for the start of vegetative growth jointly re-estimated in both the Midwest and Gulf genetic subpopulations.

**SI Dataset S4 (Dataset_4_Gulf_reproductive_growth_date.csv)**

SNP-associated effects and standard errors for the start of reproductive growth jointly re-estimated in the Gulf genetic subpopulation.

**SI Dataset S5 (Dataset_5_Midwest_reproductive_growth_date.csv)**

SNP-associated effects and standard errors for the start of reproductive growth jointly re-estimated in the Midwest genetic subpopulation.

**SI Dataset S6 (Dataset_6_Both_subpopulations_reproductive_growth_date.csv)**

SNP-associated effects and standard errors for the start of reproductive growth jointly re-estimated in both the Midwest and Gulf genetic subpopulations.

**SI Dataset S7 (Dataset_7_QTL_intervals_with_functionally_validated_homologs.csv)**

Genes in the quantitative trait loci (QTL) regions shown in Figure 4 that have functionally validated homologs in rice for vegetative or reproductive development. The first 9 columns of this table provide QTL information, the remainder provide annotation information for the switchgrass genes in the QTL interval with homologs in A. thaliana or rice that have functional validation in flowering or vegetative-growth related traits. Columns 20-30 provide the titles of papers providing functional validation of the rice homolog of the switchgrass gene.

**SI Dataset S8 (Dataset_8_QTL_mash_overlap.csv)**

Overlap between quantitative trait loci (QTL) for the start of vegetative or reproductive growth with significant mash effect estimates (with a log10(Bayes Factor) $> 2$) found using diversity panels of the Midwest, Gulf, and Both genetic subpopulations. The first five columns of this table provide QTL information, and columns 6-10 provide mash Marker information for markers within the QTL regions.
